# Multi-material delivery strategies to create deferoxamine-functionalized collagen scaffolds for craniofacial bone repair

**DOI:** 10.64898/2026.11.07.765092

**Authors:** Génesis Ríos Adorno, Ashwin Bale, Chitra Subramanian, Rohit Bhargava, Mark S. Cohen, Brendan A.C. Harley

## Abstract

Craniomaxillofacial (CMF) bone injuries are common and can occur secondary to trauma, bone resection, and congenital abnormalities. While the current standard of care remains allograft or autograft, tissue engineering approaches offer the opportunity for alternative regenerative solutions. Biomaterial design has been central in this effort, with a primary consideration being the incorporation of motifs into biomaterials that accelerate osteogenic differentiation of progenitor cells and subsequent mineral synthesis. However, angiogenic potency, and subsequent angiogenic-osteogenic coupling, represent a critical challenge, as poor angiogenic activity can impair regenerative healing. Deferoxamine (DFO) can mimic cellular hypoxia via iron chelation and has been used as an angiogenic factor. However, its size (<1 kDa) severely limits its retention in the wound area. Here, we report a multi-material approach to improve angiogenic coupling in an osteogenic biomaterial. We report methods to retain DFO in mineralized collagen scaffold via its binding to a second polymer, gelatin, that can be incorporated into the scaffold to boost angiogenic potency while maintaining its osteo-regenerative properties. We employed carbodiimide chemistry to attach DFO to a gelatin backbone and retained 79.6% of the drug over a 7-day culture. Using a composite drug-delivery approach that combines freely soluble DFO with gelatin-bound DFO into the mineralized collagen scaffold, drove increased mesenchymal stem cell metabolic activity and enhanced expression of angiogenic genes in culture over 14 days. This work provides a novel approach to advance acellular regenerative biomaterials for osteogenic-angiogenic coupling processes.

## 1. Introduction

Craniomaxillofacial (CMF) injuries account for 130,000 injuries per year in the US in civilian populations as well as more than a quarter of all battlefield injuries in recent conflicts [1, 2]. These injuries arise from trauma, bone resections, and congenital abnormalities. In critical-sized defects, the missing bone cannot regenerate, and implants are required to fill the bone gap [3]. The size of a critical-sized craniofacial defect depends on animal species, age, and injury location; studies suggest that 2 mm in mice, 14 mm in rabbits, and 25 mm in swine are large enough to impede tissue formation without intervention [4–11]. Bone autografts and allografts are the current gold-standard for care [12]. Autografts are harvested from a patient’s own body, usually from the iliac crest, but these are limited by the amount of required bone and secondary site morbidity [12]. While there is greater availability for allografts (bone tissue harvested from a cadaveric donor); risk of transplant rejection and lower osteogenic potency after sterilization has been observed [13, 14]. Regenerative biomaterials offer an alternative approach to address the scale and patient variability of CMF bone defects.

A wide range of biomaterial approaches have been described for bone tissue engineering, with many containing similar motifs of the combination of inorganic and organic materials (*e.g.*, collagen, mineral) [3, 15]. We have previously reported a freeze-drying approach to fabricate a mineralized collagen scaffold that contains an open-pore architecture (90% porosity, 120 µm diameter) that facilitates cell infiltration and have optimized its composition and mechanical properties to promote osteogenic differentiation and mineral formation without the need for conventional exogenous factors (e.g., BMP-2; osteogenic media) [10, 16–19]. We have also reported approaches to tune its glycosaminoglycan content, pore shape, and biomolecule retention to aid the secretion or expression of bone-related markers such as ALP, OPG, VEGF, and BMP-2 [20–23]. However, angiogenesis and osteogenesis are intimately coupled, and a predisposing factor for failed healing occurs with impaired angiogenesis at the defect-implant margin [24]. Poor vascular healing is also the major challenge in precision medicine, with disparate angiogenic activity seen as a result of patient age (e.g., increased senescence [25]) or radiation with therapy (e.g., cancer) [26, 27].

The first two weeks after implantation are critical for vascular recruitment, the coordination of angiogenic and osteogenic activity, and eventual regeneration [28–36]. Endothelial cell recruitment is required to form vascular tissue and facilitate nutrient and oxygen transport in the wound area [3, 28]. While the osteogenic and angiogenic coupling in flat bone healing is not completely understood, with some studies suggesting they are independent processes [37], the scale of bone formation required for fracture or defect healing necessitates metabolic support that arises from efficient vasculature formation, motivating efforts to develop biomaterials that can support osteogenic and angiogenic processes synergistically [38, 39]. Pro-angiogenic growth factors (VEGF, PDGF and TGFβ families) have been incorporated into biomaterials in a variety of formats to stimulate endothelial cell recruitment and vascular formation. These approaches face challenges from poor growth factor (GF) retention (not enough factor) or unintended effects of often supra-physiological GF doses [39]. Other approaches have included pre-vascularization steps using endothelial cell cultures in vitro; however, clinical translation of such approaches would require extensive pre-operative cell culture, adding barriers into clinical translation applications and regulatory compliance [37, 40–42].

Deferoxamine (DFO), an FDA-approved iron chelator, has been used for a wide range of biomedical applications, including wound healing, diabetes, ischemia, bone nonunion, and cancer treatment, due to its ability to promote vascularization and down regulation of inflammatory responses [38, 43, 44]. HIF-1α regulates angiogenic pathways, such as the upregulation of VEGF and SDF-1 to promote endogenous cell recruitment and vascular formation. However, this can only happen under hypoxic conditions, as HIF-1α degrades in the presence of oxygen. DFO can stabilize HIF-1α under normoxic conditions by preventing HIF prolyl hydroxylase domain protein from degrading HIF-1α to prolong angiogenic effects, making it extremely promising for biomedical applications [45, 46]. However, its small size (~ 600 Da) limits its retention at the injury site and contributes to a shorter half-life of 20-30 minutes, resulting in rapid systemic clearance [38, 47]. Recent approaches have explored strategies such as nanoparticle/microsphere in hydrogel system to prolong DFO retention in the wound site. Recently, *Donneys et al.* described an approach to conjugate the primary free amine in DFO to the carboxyl group in hyaluronic acid (HA). The resulted HA-DFO composite prolonged DFO retention for over two weeks while providing an osteoconductive and anti-inflammatory matrix that enhanced neovascularization and bone healing in an irradiated rat bone nonunion model [38]. While effective, this strategy relied entirely on the angiogenic activity of HA-DFO without considering design motifs to simultaneously promote osteogenic activity. Separately, we recently reported a strategy to create a library of gelatin hydrogel formulations whose rheological and mechanical properties could be tailored to facilitate their inclusion into a porous collagen scaffold [48], suggesting a multi-material composite approach to combine the osteogenic nature of a mineralized collagen scaffold with the DFO-retention capacity of a hydrogel carrier.

Here we describe an effort to enhance the angiogenic properties of a porous mineralized collagen scaffold via the inclusion of DFO while retaining its osteogenic properties. Angiogenesis occurs early in the healing process (approx. 7 days after callus formation). Therefore, there is a critical window for implantation success, and controlled retention of DFO is desired during this time frame to avoid burst release immediately after implantation. Inclusion of materials with different diffusion coefficients might help control the release of molecules, with hydrogels being extensively used as injectable drug carriers with tunable release kinetics defined by polymer size, crosslink chemistry, and environmental conditions [49–51]. Previously, we outlined the synthesis of maleimide-functionalized gelatin (GelMAL) materials to form stable gels [52]. Here, we describe bioconjugation of DFO to GelMAL as a drug-delivery carrier as well as the design of a DFO-collagen composite scaffold as an osteo-angiogenic-inducer biomaterial. We quantify DFO retention in a gelatin hydrogel carrier, HIF-1α stabilization in response to extended DFO exposure, and VEGF upregulation in response to rapid DFO availability without affecting scaffold osteogenic properties.

## 2. Materials and Methods

### 2.1. Synthesis of Gelatin Maleimide (GelMAL)

Gelatin maleimide was synthesized as described previously [52]. Briefly, 10 mL 0.5 M 2-Morpholinoethanesulphonic acid buffer (MES, pH 6.0) and 8 mL dimethyl sulfoxide (DMSO) were used to dissolve 0.5 g of Type B Bovine-derived Gelatin (Sigma Aldrich, MO) at 40°C. Later, 0.24 g of succinimidyl 3-maleimidopropionate (Tokyo Chemical Industry Co., OR) was dissolved in 2 mL of DMSO and added to the reaction. The reaction ran for 24 h at 40 °C and 720 RPM after the pH was adjusted to 4.5. This was followed by 5 days of dialysis against acidic water (pH 3.25) at 37°C, and 7 days of lyophilization. The material was stored at −20°C until use.

### 2.2. Synthesis of GelMAL-DFO

Functionalization of GelMAL with DFO was achieved using carbodiimide chemistry with 1-Ethyl-3-[3-dimethylaminopropyl]carbodiimide hydrochloride (EDC, Oakwood Chemical, SC) and NHS (N-Hydroxysuccinimide, Sigma Aldrich) [38]. 0.5g of GelMAL was dissolved in 0.5 M MES buffer (pH 4.5) at 37 °C and 720 RPM. Once dissolved, 197 mg of DFO were dissolved in 4 mL 0.5 M MES buffer and added to the mixture. EDC and NHS were dissolved in reaction buffer and added into the reaction vial at a 1:1:1 COOH: EDC: NHS molar ratio. The pH of the reaction was adjusted to 4.5, and the reaction was run for 24 h at 37 °C. After reaction completion, the solution was dialyzed for 3 days in acidic water (pH 3.25), frozen overnight, and lyophilized for 7 days. The dried material was stored at −20°C until use.

### 2.3. NMR characterization

Successful functionalization of maleimides and deferoxamine was validated via Nuclear Magnetic Resonance (H-NMR) as previously demonstrated [52]. Briefly, NMR spectra of 12.5 mg/mL gelatin, GelMAL or GelMAL-DFO solution in D_2_O were collected at 25 °C in an Agilent 600 MHz NMR spectrometer (64 scans, 5s spin, Santa Clara, CA), and phase and baseline corrected. The chemical shift scale was adjusted to the residual solvent peak (4.79 ppm). The degree of functionalization of maleimide peaks was determined relative to the phenylalanine peak.

### 2.4. FTIR characterization

Fourier Transform Infrared spectroscopic (FT-IR) data from gelatin, GelMAL, and GelMAL-DFO macromers were measured using attenuated total reflectance (ATR) sampling using a diamond crystal. Full spectra were acquired across the mid-IR range using an undersampling ratio of 4 (0-3950 cm^−1^) and truncated to a range of 900–3300 cm^−1^ at a spectral resolution of 4 cm^−1^. The raw spectra were normalized and its second derivative vector-normalized, as previously described [53, 54], for comparison between conditions.

### 2.5. Quantification of DFO within gelatin hydrogels

The concentration of DFO in GelMAL-DFO was determined using a FeCl_3_ colorimetric assay and adapted from previously outlined methods [38]. 10 mg/mL of GelMAL-DFO was first digested overnight at 37°C in 100 U/mL collagenase II (ThermoFisher Scientific) and 3 mM of CaCl_2_ in Hank’s Balanced Salt Solution (HBSS, Corning, NY), GelMAL material was used as a control. 500 µL of digested material or deferoxamine mesylate standard solution (10 – 0.02 mg/mL) was mixed with 500 µL of 6 mM FeCl_3_ solution in HBSS, placed in a shaker, and incubated overnight at room temperature. Absorbance was measured at 430 nm in triplicate using a Tecan Infinite M200 Plate Reader (Tecan, Männerdorf, Switzerland).

### 2.6. Hydrogel fabrication and sterilization

Hydrogel fabrication was adapted from previously reported methods [52]. GelMAL or GelMAL-DFO were UV-sterilized for 30 minutes and stored at −20 °C until use. 4 wt.% GelMAL or GelMAL-DFO were dissolved in sterile PBS at 37 °C. For different concentrations of DFO, GelMAL and GelMAL-DFO were mixed prior to material dissolution. If cells were involved in the experiment, cell suspension was added at this time. 50 µL of gel precursor was then transferred into a 5 mm-diameter Teflon mold and gelled at 4 °C for 30 min. Next, 1 mL of 0.13 mg/mL chilled Dithiothreitol (DTT, TCI America, Tokyo) was added, and the gels were incubated for 40 min at 4 °C to complete crosslinking via thiol-Michael Addition. Gels were washed in PBS twice after crosslink. Basal cell media (low glucose Dulbecco’s Modified Eagle Medium with added glutamine (School of Chemical Sciences Cell Media Facility, University of Illinois Urbana-Champaign), 10% mesenchymal stem cell fetal bovine serum (Gemini Bio Products, West Sacramento, CA), and 1% antibiotic-antimycotic (Gibco, Billings, MA)) was added to cell-seeded gels and incubated at 37 °C and 5% CO_2_; otherwise, the gels were incubated in PBS.

### 2.7. DFO release quantification

4 wt.% GelMAL-DFO gels were fabricated as described in the previous section. Gels were transferred into PBS and incubated at 37 °C. PBS was collected every day for 7 days after fabrication. PBS media was stored at −20 °C until quantification. 24 h before quantification, samples were fridge thaw, and 500 µL were transferred into vials. Deferoxamine mesylate was used to prepare 2-fold serial dilution standard solutions (10 – 0.02 mg/mL). DFO analytical quantification was determined using Liquid Chromatography-Mass Spectrometry (LCMS) in the Mass Spectrometry Lab at the School of Chemical Sciences at the University of Illinois Urbana-Champaign [55].

### 2.8. Mineralized collagen scaffold fabrication

#### 2.8.1. Scaffold fabrication

Mineralized collagen scaffold was fabricated as described previously [19, 20, 23]. Briefly, a collagen suspension was fabricated by homogenizing 1.9 w/v% type I bovine Collagen (Collagen Matrix Inc., NJ), 0.84 w/v% Chondroitin-6-sulfate (Spectrum Chemical Mfg. Corp., CA) and calcium salts (0.9 w/v% CaOH and 0.4 w/v.% Ca (NO_3_)_2_•4H_2_O, Sigma-Aldrich, St. Louis, MO) in 0.1456 M phosphoric acid/0.037 M CaOH buffer. Suspension was stored overnight at 4 °C and degassed prior to lyophilization. Collagen slurry was loaded into an aluminum mold and freeze dried at 1 °C/min to −10 °C, followed by a 2 h hold step at −10 °C and sublimated at 0 °C and 0.2 Torr using a Genesis freeze-dryer (VirTis, NY, USA) to form an isotropic structure. Collagen sheets were cut using a 6 mm biopsy punch.

#### 2.8.2. Sterilization, hydration, and crosslinking

Collagen scaffolds were sterilized using an AN74i Anprolene gas sterilizer (Andersen Sterilizers Inc, Haw River, NC), through a 12 h cycle of ethylene oxide treatment as previously described [56, 57]. Materials were hydrated and crosslinked following previously outlined methods [20, 58]. All steps were performed under sterile conditions at room temperature. Briefly, scaffolds were soaked in 100% ethanol for 2 h and washed in PBS twice for 30 minutes each before treatment with an EDC/NHS (5:2:1, EDC: NHS: COOH molar ratio) solution for 1.5 h. Finally, the scaffolds were washed twice with PBS for 30 min and incubated at 37 °C and 5% CO_2_ in cell culture media for 48 h prior to cell seeding.

### 2.9. Collagen-gelatin Composite Fabrication

A composite scaffold comprising a collagen scaffold and GelMAL or GelMAL-DFO was fabricated by combining both materials. 4 wt.% GelMAL or GelMAL-DFO hydrogel precursor was dissolved in PBS, and 25 µL was pipetted on top of the hydrated collagen scaffold. The hydrogel component in the composite material was crosslinked as described in the previous section [52]. Composites were gelled at 4 °C for 30 min. The concentration of bound DFO in the composite (DFO or DFO + Free DFO groups) was 200 µM relative to 110 µL of material. Then, 1 mL of chilled DTT was used to crosslink the GelMAL or GelMAL-DFO for 40 minutes at 4 °C. Finally, the composites were washed with PBS. Cells were seeded following the collagen scaffold seeding strategy as discussed in the following subsection. For Free DFO and DFO + Free DFO groups, media was supplemented with 20 µM of DFO (1 mL/ scaffold).

### 2.10. Human mesenchymal stem cell culture

Adipose-derived human mesenchymal stem cells (hMSCs, Male, Caucasian, Essent Biologics, Lot no. 206079) were cultured at 37 °C and 5% CO_2_ in RoosterNourish-MSC (RoosterBio, MD) expansion media until passage 5. hMSCs were lifted and seeded at 150,000 cells onto collagen scaffolds or composites as described previously [22]. Briefly, 5 µL containing 75,000 hMSCs was seeded on one side of the scaffold and incubated at 37 °C for 30 min. Scaffolds were then flipped, and an additional 5 µL of cell suspension was added (total of 150,000 cells/scaffold) and incubated for 1.5 h at 37 °C for cell attachment. Following the incubation time, 1 mL of cell media was added. For hydrogel-based experiments, a cell aliquot containing 150,000 cells was added to the hydrogel precursor before the mixture was transferred into the Teflon molds. Cell media was changed every 3 days until experiment completion.

### 2.11. Metabolic Activity and DNA quantification

Cell number (n=6) was directly quantified from isolated DNA using a DNAeasy Blood and Tissue kit (Qiagen, MD) on experimental days 1,3,7, and 14 as described on previous publications [59]. The DNA was eluted at 200 µL, and prior to isolation, samples were cut into 8 pieces and incubated at 56 °C overnight on lysis buffer. Known values of cell number were used to generate a calibration curve on experimental Day 0 following manufacturer’s protocol for cell suspensions to quantify isolated DNA. DNA concentration was quantified in triplicate via Nanodrop One Spectrophotometer (Thermo Scientific).

Metabolic activity was measured at experimental days 1,3,7, and 14 using an AlamarBlue assay (n=6, Invitrogen, Carlsbad, CA) [60, 61]. Samples (scaffolds, or hydrogels) were incubated in a 9:1 volume ratio of basal media to AlamarBlue solution for 90 min at 37 °C, 5% CO_2_ and 100 RPM. Samples were returned to basal media and reused for the following time points. Fluorescence of conditioned media was measured in triplicate using a F200 spectrophotometer (Tecan, Männedorf, Switzerland) at 540/580 nm. A standard curve on Day 0 was generated from cell suspension with known cell concentration.

### 2.12. RNA isolation and Gene Expression

RNA (n=6) was isolated as described in previously outlined methods [61, 62]. Samples were cut into 8 pieces and transferred into a PhaseShield Gel tube (LifeSCT, MD) and incubated in 1 mL of Trizol (Invitrogen) for at least 5 min, followed by 200 µL of chloroform at room temperature. Tubes were then centrifuged at 15,000 g for 15 min at 4 °C. The clear supernatant phase was transferred into a clean centrifuge tube and mixed with an equal volume of ethanol. RNA purification was performed using a Zymo RNA Clean and Concentrator-5 kit (Zymo, CA) following the manufacturer’s protocol, performing centrifugation steps at 15,000 g for 30 s and a 15 µL elution step. RNA concentration was measured in a NanoDrop One Spectrophotometer. Synthesis of cDNA from RNA templates was performed using a QuantiTect Reverse Transcription kit (Qiagen) and a S100 thermal cycler (Rio-Rad, CA, USA) as described before [61, 62]. Real-time quantitative PCR (RT-qPCR) was performed using 50 ng cDNA, TaqMan™ primers (Table 1), and TaqMan™ Fast Advanced Master Mix (Applied Systems, CA, USA). Reactions (10 µL) were run in duplicates in a QuantStudio 7 Flex Real Time PCR System (ThermoFisher Scientific). Data analysis followed the ΔΔCt method, normalizing *GAPDH* as the housekeeping gene and Day 0 gene expression as the control group.

**Table 1.** Taqman Gene Primers for RT-qPCR.

| **Gene Name** | **Gene Symbol** | **Taqman Assay ID** |
|---|---|---|
| Glyceraldehyde-3-phosphate dehydrogenase | GAPDH | HS02786624_G1 |
| Hypoxia inducible factor 1 alpha subunit | HIF1a | HS00153153_M1 |
| Vascular endothelial growth factor A | VEGF | HS00900055_M1 |

### 2.13. Secretome Analysis

Protein release (OPG, VEGF, n=3) was measured using human DuoSet ELISA kits (R&D Systems, MN, USA, cat no. DY805, and DY293B, respectively) [20, 22]. Media from cultured samples were collected every three days and stored at −20°C until the assay was conducted. Assays were performed as described in kit protocols. Samples (10 for OPG and 12.5 µL VEGF) were diluted in reagent diluent right before the sample incubation step, and absorbance was measured utilizing a M200 spectrometer Tecan plate reader (Tecan) in duplicate. Data was analyzed using a 2^nd^ order polynomial regression from a generated calibration curve.

### 2.14. Vascular Analysis

Matrigel tubule formation was performed as described [20]. Human Umbilical Vein Endothelial cells passage 4 (HUVECs, Caucasian, Lonza, Lot no. 18TL05850) were cultured at 37 °C, and 5% CO_2_ until 80% confluency (around 5 days) in Endothelial cell basal medium-2 (Lonza, Cat no. CC-3156). A 150 µL pooled volume of conditioned media (Day 3 or 14) from the scaffold composite experiment was mixed with 10,000 HUVECs (n=6) and transferred into a Phenol Red-Free Corning™ Matrigel™ Matrix-3D plate (Corning). Wells were imaged using the Leica DMI8 Confocal microscope at 0, 6 and 12 h post cell seeding. The center area of the well was imaged throughout the experiment, and Brightfield images were analyzed using the “Angiogenesis Analyzer” macros tool in ImageJ (NIH, Maryland, USA) to quantify tube and junction formation [20, 63]. The average branch length was calculated from total branch length by the total number of branches.

### 2.15. Statistics and Data Visualization

Statistical analysis was performed on R (4.6.0) via RStudio (2026.04.0+ 526). Raw data were evaluated for outliers using a Grubbs test in OriginPro 2025b before statistical analysis. Outliers were removed from statistical analysis and visualized data. A p<0.05 was used as a criterion to reject the null hypothesis on all tests performed. After fitting the dataset to a linear model, normal distribution was tested using the Shapiro-Wilk test and checked for equal variance with the Levene test. ANOVA and Tukey’s post-hoc tests were used for data with normal distribution and equal variance. Kruskal-Wallis and Dunn tests with Benjamini-Hochberg correction were used if the data were non-normal and variances were unequal. Normal distribution with unequal variances was tested using Welch’s ANOVA test with Games-Howell post-hoc. Data with neither normal distribution nor equal variances were tested via Welch’s Heteroscedastic F test with trimmed means and winsorized variances and a Games-Howell post-hoc test. Data sets with 2 groups were tested following a t-test if they were normally distributed with equal variances or a Welch’s t-test if data were normally distributed, but the variances were unequal.

Conceptual illustrations were created using BioRender. Data Figures were created using OriginPro 2025b. Box-and-whisker plots denote interquartile range (IQR, box) ± 1.5 outlier (whiskers), with the straight line indicating the median and the ‘x’ or ‘□’ symbol the mean. Bar plots represent the mean ± 1 standard deviation, with the straight line indicating the median. Similarly, line plots represent the average value ± 1 standard deviation. Statistical symbols on the plots denote significance between groups, *p<0.05, **p<0.01, ***p<0.001, **** p<0.0001 and n.s. p>0.05. Groups with different letters in data plots denote statistical significance (p <0.05).

## 3. Results

### 3.1. DFO supplemented in media promotes the upregulation of VEGF in hMSCs

We first evaluated the effect of a range of soluble DFO concentrations on hMSC seeded mineralized collagen scaffolds (Fig. 1A). DFO was added with basal cell culture media on culture Day 0 (after hMSCs were seeded into the scaffolds), with the initial media replaced on Day 3 and no further DFO supplementation through the end of the 14-day experiment. High concentrations of DFO (≥ 50 µM) led to reduced overall numbers of hMSCs proliferation but increased the metabolic activity of the remaining cells (Fig. S1A-B). Analysis of hMSC gene expression patterns (Fig. 1B-C) revealed a rapid, dose-dependent effect of increasing DFO concentration on increasing VEGF expression, but only within the first 3 days after DFO exposure. Analysis of HIF-1α expression revealed downregulation at the highest levels (50 and 100 µM) of DFO exposure at day 3, followed by DFO dose-dependent increases in HIF-1α at later times (day 14). The mineralized collagen scaffold has been previously shown to promote hMSC secretion of VEGF and OPG, a glycoprotein inhibitor of osteoclast activity; while exposure to a single dose of soluble DFO increased early (day 3) VEGF production, high levels of DFO exposure led to reduced overall secretion of OPG and VEGF through extended (up to 14 day) culture (Fig. S1C-D).

**Figure 1.**
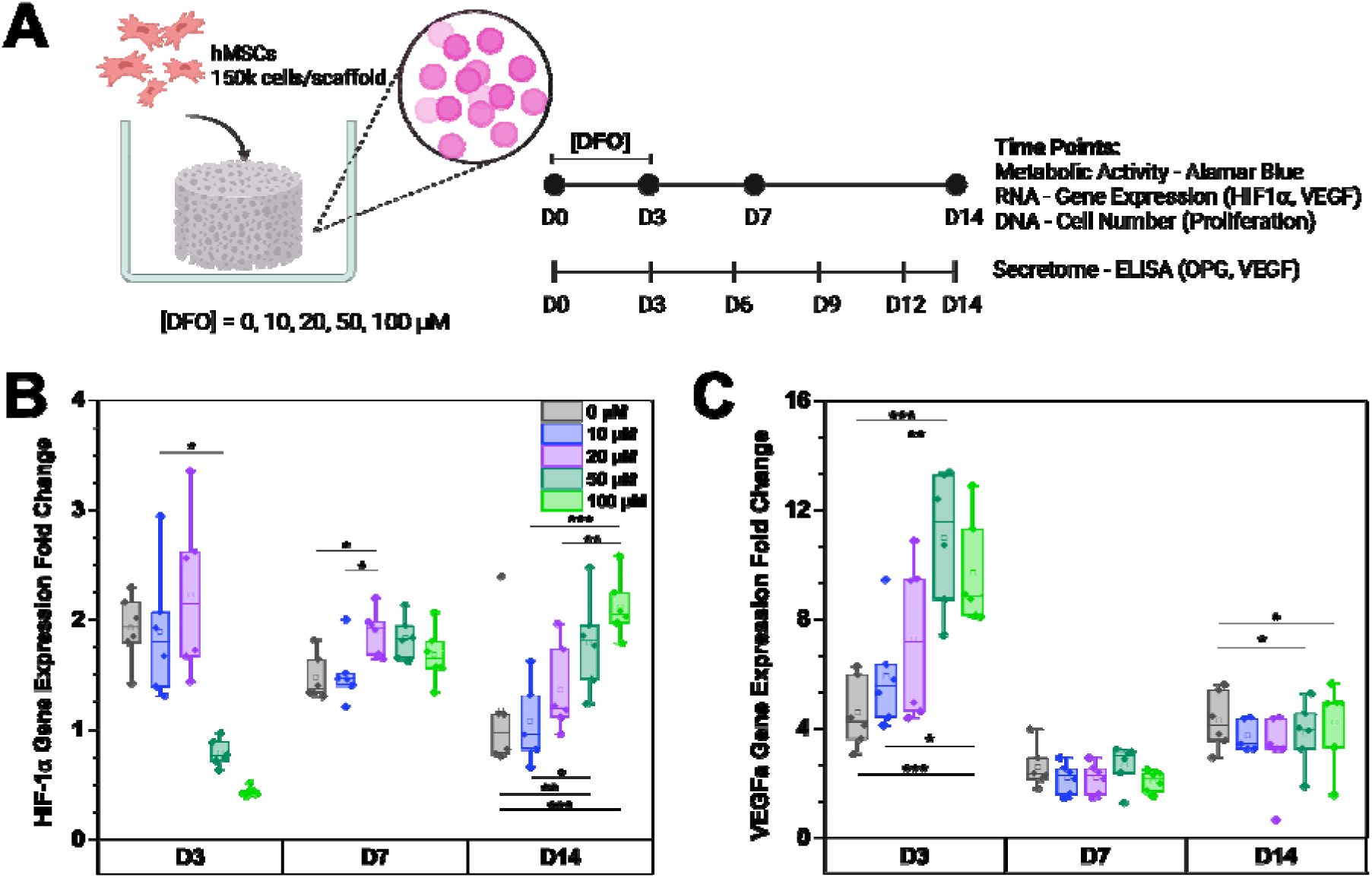
DFO supplemented media effects on hMSCs. **A.** Experimental Outline. Multiple DFO concentrations were supplemented in the media as a single dose on experimental day 0 and removed during the first media change after 3 days of culture. **B.** Box plot of HIF-1α gene expression fold changes across days of culture. **C.** Box plot of VEGF gene expression fold changes across days of culture. Gene expression (n=6) was quantified through RT-qPCR and analyzed following ΔΔC_T_ method with GAPDH and Day 0 as controls. Box plot diagrams show IQR (box) ± 1.5 outliers (whiskers) and mean (□). * p<0.05, ** p<0.01, and *** p <0.001.

### 3.2. DFO can be conjugated to MAL and COOH groups in gelatin backbone

We then examined strategies to retain DFO within the mineralized collagen scaffold via conjugation to a gelatin backbone (Fig. 2A). First, a maleimide-functionalized gelatin (GelMAL) macromer was created via exposure to N-succinimidyl 3-Maleimidopropionate. EDC/NHS carbodiimide chemistry was used to attach the free primary amine in DFO to either COOH or maleimide (MAL) groups on the GelMAL. Successful conjugation was confirmed via 1H-NMR, with the characteristic MAL peak (6.82 ppm; versus phenylalanine) used to calculate degree of gelatin functionalization (DoF), finding MAL DoF of 61% in unmodified GelMAL but only 30% for GelMAL-DFO (Fig. 2B). NMR spectra also show several peaks across the 2.9 to 1.25 ppm range corresponding to DFO. Peaks at 2.86 and 2.75 ppm confirm gelatin-bound DFO rather than free DFO molecules in the sample [38, 64].

**Figure 2.**
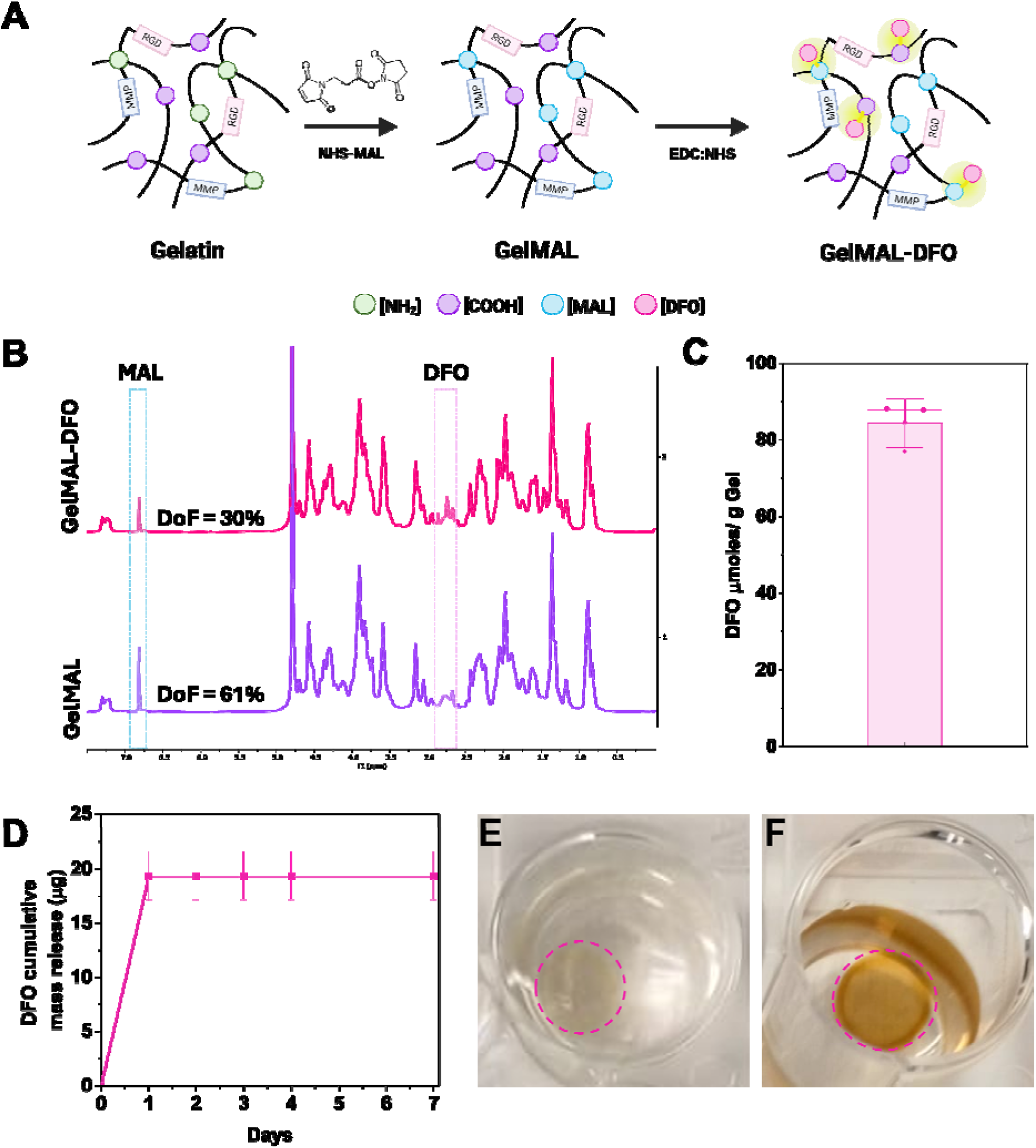
DFO-gelatin chemical binding and characterization. **A**. Native gelatin was functionalized with maleimides using N-succinimidyl 3-Maleimidopropionate (GelMAL). Subsequently, primary free amine in DFO was conjugated to carboxyls and maleimides in gelatin following carbodiimide chemistry (GelMAL-DFO). **B**. 1H-NMR spectra of GelMAL and GelMAL-DFO. Maleimide addition was confirmed by a new peak at 6.82 ppm. DFO attachment generated a reduction in the maleimide peak, and peaks forming at 2.86 and 2.75 ppm corresponding to DFO characteristic peaks. **C**. DFO concentration on functionalized gelatin. **D**. DFO cumulative release from functionalized 4 wt.% GelMAL-DFO for 7 days in PBS at 37 °C. **E**. 4 wt.% GelMAL-DFO hydrogel in 3mM FeCl_3_ at t=0 and **F**. At t=16 h of FeCl_3_ exposure. Plots represent the average value ± SD.

ATR-FTIR spectroscopy was also used to characterize synthesized materials by examining several peaks of interest (Fig. S2A-D). The Amide I region, which arises predominantly from the C=O stretching vibration of the peptide backbone, showed a reproducible difference between unmodified gelatin and the functionalized materials. In gelatin, the second derivative resolved a minimum near 1621 cm^−1^, a position associated with intermolecular hydrogen-bonded, aggregated backbone environments [65]. In both GelMAL and GelMAL-DFO this component was shifted to approximately 1629 cm^−1^ (Fig. S2C). A second region of interest, 1700–1750 cm^−1^, contains carbonyl stretching vibrations of the maleimide group. N-substituted maleimides characteristically display an asymmetric C=O stretch near 1700-1720 cm^−1^ [66]. Consistent with this vibrational mode, a distinct second-derivative minimum was observed at 1710 cm^−1^ in GelMAL and GelMAL-DFO that was absent in unmodified gelatin (Fig. S2B), supporting successful maleimide functionalization. The 1000–1100 cm^−1^ region was additionally examined as a possible corroborating marker of DFO incorporation. DFO contains three hydroxamic acid (hydroxamate) groups responsible for its iron-chelating activity; while the hydroxamate N–O stretch is most consistently reported near 900–960 cm^−1^, some studies have also attributed features extending into the 1000–1100 cm^−1^ region to hydroxamate- and amine-associated vibrations [67–69]. In our analysis, increases in second-derivative intensity were observed at 1014, 1051, and 1061 cm^−1^ in the DFO-loaded samples (Fig. S2D). This region was therefore included as a secondary line of evidence, in addition to the carbonyl-region analysis described above.

The functional analysis of conjugated DFO was performed via an Iron assay, in which DFO forms a complex with iron, resulting in an orange/red color upon chelation. GelMAL-DFO macromer was digested with Collagenase II and CaCl_2_, then mixed with 3 mM FeCl_2_ for 16 h. GelMAL was used as a negative control, and results were compared to a DFO calibration curve. This analysis revealed this conjugation created a GelMAL-DFO with 47.3 ± 3.6 mg DFO/g Gelatin (84.3 ± 6.4 µmoles DFO/g Gelatin; Fig. 2C). Finally, DFO release from GelMAL-DFO hydrogels was quantified via liquid chromatography mass spectroscopy over 7 days. GelMAL-DFO gels released 19.3 ± 2.2 µg of DFO in the first 24 h, with no additional DFO detected after 24 h (Fig. 2D), suggesting the GelMAL-DFO macromer is able to retain 79.6% of loaded DFO over 7 days. To confirm DFO retention in the GelMAL-DFO conjugate, after 48 hours of incubation, the GelMAL-DFO hydrogel was exposed to 3 mM FeCl_2_ to visualize the presence of unreleased DFO in the gel (Fig. 2E). GelMAL-DFO gels showed an intense orange color, implying DFO availability (Fig. 2F).

### 3.3. Conjugated DFO increases hMSCs VEGF secretion

We then evaluated the ability for DFO-conjugated GelMAL hydrogels to enhance the angiogenic activity of hMSCs. Here, 150,000 cells were encapsulated in 4 wt.% GelMAL-DFO hydrogels containing a range of DFO concentrations (0 - 1000µM), then maintained in culture for up to 14 days (Fig. 3A). DFO concentrations were achieved by mixing mass ratios of GelMAL to GelMAL-DFO macromers. Cell number was quantified as DNA yields normalized to 0 µM DFO hydrogel (GelMAL) on day 1, showing some reduction in cell numbers in the presence of high concentration (>200 µM DFO) of DFO (Fig. 3B). Similarly, DFO-functionalized hydrogels reduced cell metabolic activity at early stages but the effect was not significant after day 7 (Fig. 3C). Interestingly, hMSCs showed increased secretion of VEGF in DFO-functionalized hydrogels across the 14 days culture experiment, with a significant (p < 0.05) effect seen for 200 µM GelMAL-DFO hydrogels by day 12.

**Figure 3.**
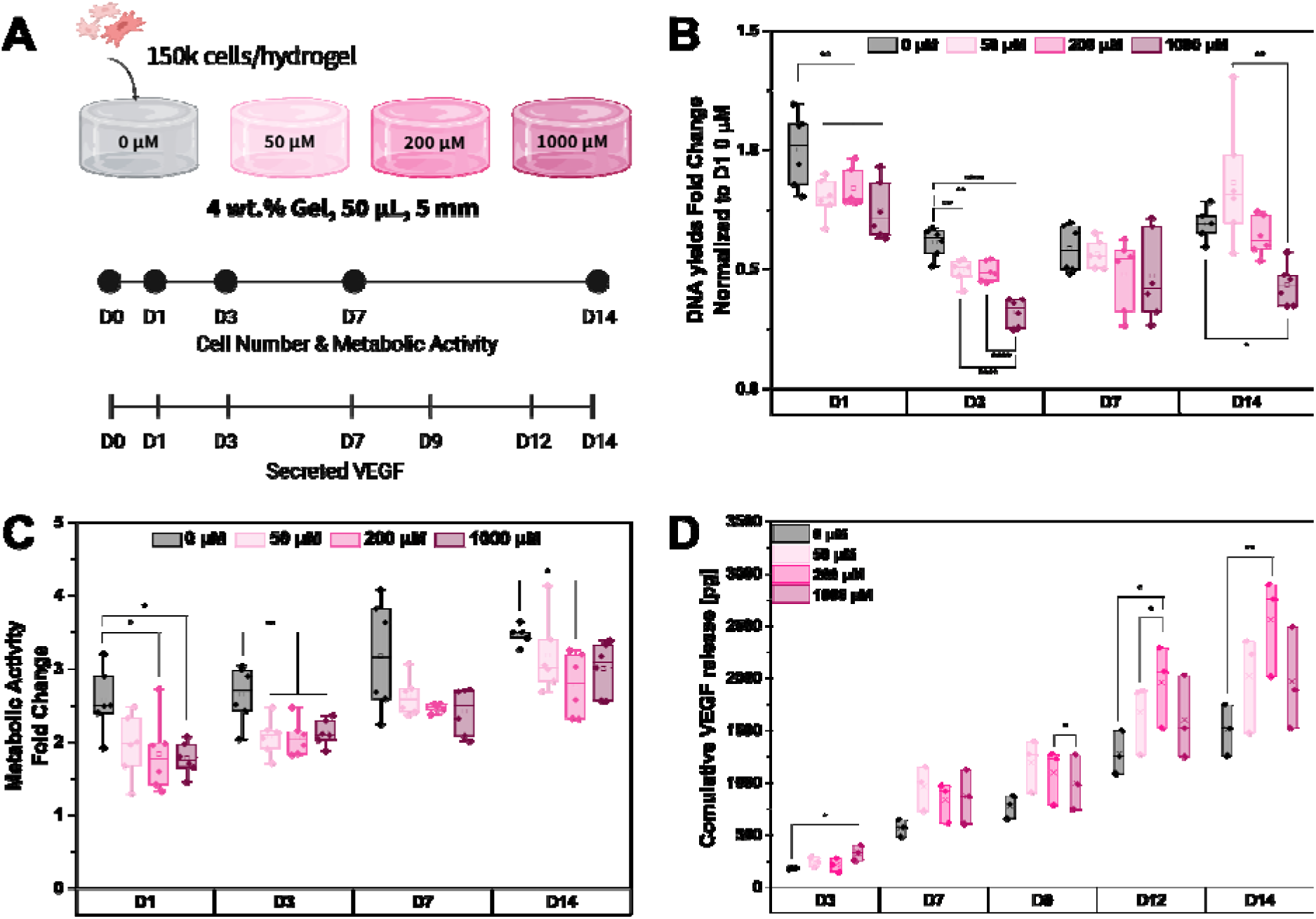
GelMAL-DFO effects on hMSCs. **A**. 3 x10^6^ hMSCs/mL were encapsulated in 4 wt.% GelMAL, or GelMAL-DFO mix hydrogel precursors and cultured for 14 days. DNA, metabolic activity and VEGF secretion were measured at several timepoints. **B**. Normalized DNA yields to D1 0 µM DFO across time. **C**. Metabolic activity fold changes across time. **D**. VEGF cumulative release. Box plots diagrams show IQR (box) ± 1.5 outliers (whiskers) and mean (□, or x). * p<0.05, ** p<0.01, and **** p <0.0001.

### 3.4. Gel-DFO containing scaffolds show a slower rate of cell proliferation but increased cell metabolic activity

We subsequently pursued a multi-material strategy, combining GelMAL-DFO hydrogels with the mineralized collagen scaffold seeded with hMSCs to accelerate osteogenic-angiogenic coupling. A DFO-collagen composite was fabricated by pipetting 25 µL of GelMAL or GelMAL-DFO (200µM DFO) hydrogel precursor on top of a hydrated mineralized collagen scaffold, and hydrogel crosslinking was achieved through a thiol-Michael addition reaction using DTT [52]. hMSCs were then seeded onto either a control mineralized scaffold (no hydrogel) or GelMAL-scaffold or GelMAL-DFO-scaffold composites (200µM DFO). All scaffolds were placed into basal media, with additional experimental groups of the hMSC seeded scaffold and hMSC seeded GelMAL-DFO-scaffold composite receiving a single dose of 20µM soluble DFO at the start of culture. All constructs were then cultured for 14 days (Fig. 4A). Inclusion of a DFO-containing GelMAL hydrogel into the scaffold significantly reduced cell proliferation in the first week of culture; however, no differences were observed between groups by day 14 (Fig. 4B). Notably, the composite containing hydrogel-bound and soluble DFO (Gel-DFO + free DFO) composite showed the fastest cell proliferation between day 7 and day 14. Composites containing GelMAL conjugated-DFO also showed significantly (p < 0.05) highest metabolic activity throughout the 14 days of culture (Fig. 4C). Interestingly, the increase in metabolic activity was tied to hydrogel-functionalized (not soluble) DFO, suggesting the effect is due to the sustained presentation of the DFO.

**Figure 4.**
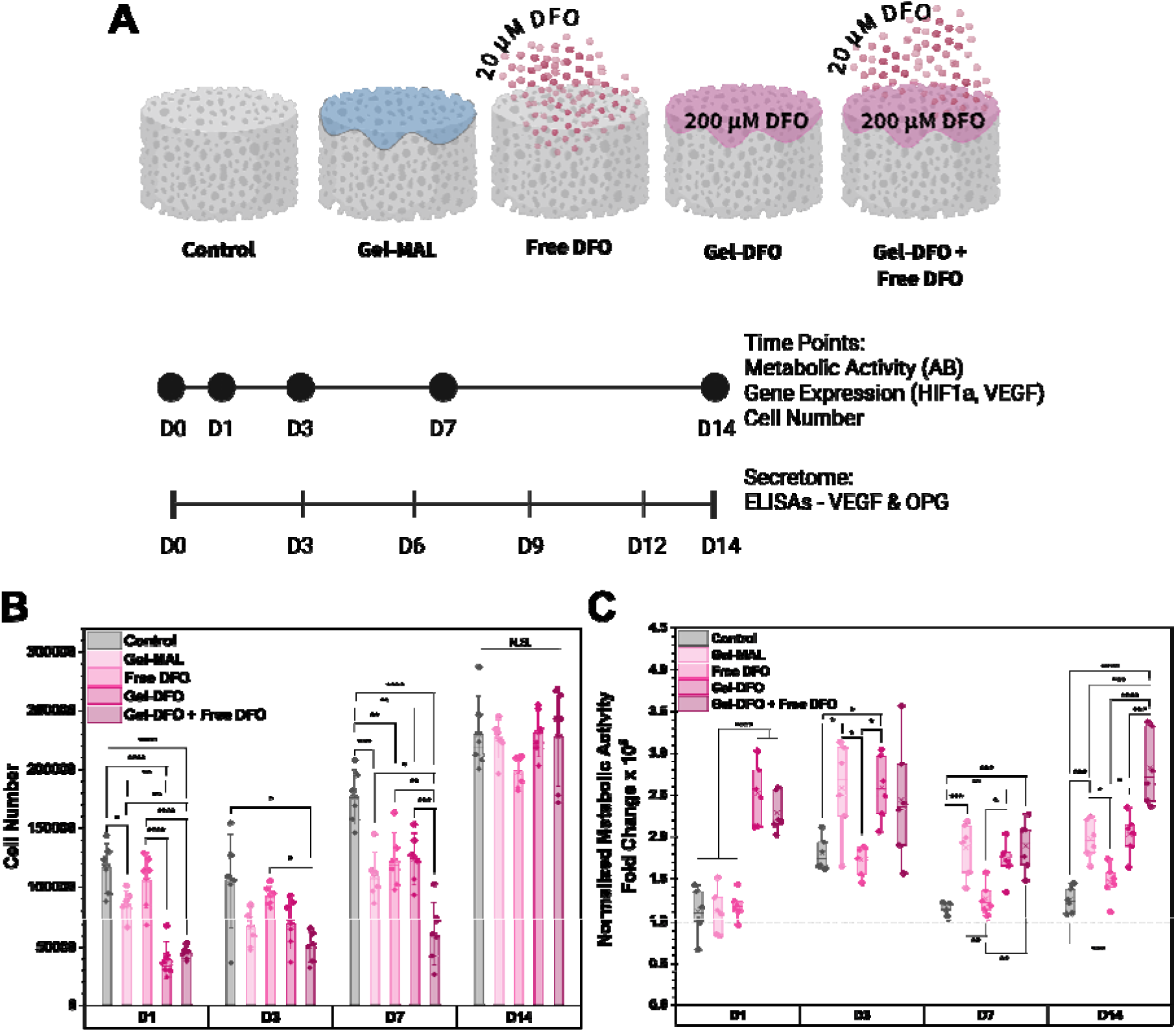
In-vitro test of DFO-Collagen materials. **A**. Composite materials are comprised of 25 µL of 4 wt.% GelMAL and/or GelMAL-DFO. The free DFO group was supplemented with 20 µM of DFO in media until the first media change at Day 3. The DFO concentration in GelMAL-DFO was 200 µM. Scaffolds were seeded with 150,000 cells on Day 0 and cultured for 14 days. Cell number, metabolic activity, gene expression and secreted factors were quantified across different time points. **B**. Cell number across days. **C**. Metabolic activity fold changes normalized to cell number. Bar plots represent the average value ± SD, and Box plots diagrams show IQR (box) ± 1.5 outliers (whiskers) and mean (x). * p<0.05, ** p<0.01, *** p<0.001, **** p <0.0001, N.S. no statistical significance.

### 3.5. Addition of DFO-functionalized hydrogel composites induced upregulation of angiogenic markers at early and late stages of in vitro culture

We subsequently examined the expression level of key angiogenic and osteogenic markers via gene expression and functional secretion. Gene expression was characterized by RT-qPCR using the ΔΔCt method, with day 0 cells as the control and GAPDH as the housekeeping gene. Notably, HIF-1α expression levels were enhanced via the orientation of soluble and hydrogel-bound DFO across the entire culture period with the effects most significant both early (day 1) and late (day 14; Fig. 5A). The presence of soluble and hydrogel-bound DFO significantly increased VEGF expression and otherwise showed an increase with time across all the conditions (Fig. 5B). Analysis of the cumulative release of VEGF and OPG into the media revealed that osteoprotegerin (OPG) secretion was not affected by either the presence of the hydrogel carrier nor the presence of soluble or hydrogel bound DFO or hydrogel inclusion (Fig. 6A). However, inclusion of soluble or hydrogel bound DFO with hMSC-seeded mineralized collagen scaffolds led to a significant reduction in total VEGF secretion across the 14 day culture (Fig. 6B).

**Figure 5.**
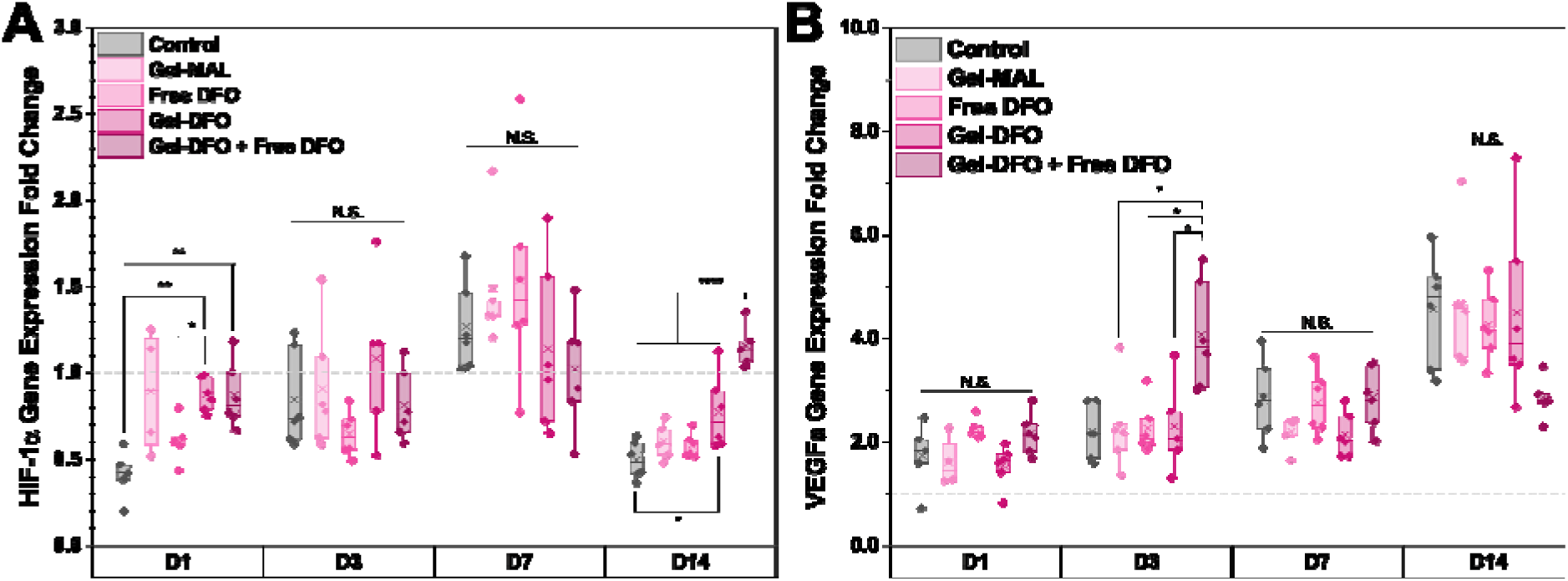
Gene expression characterization. **A**. Hif-1α expression fold change. **B**. VEGFa expression fold change. Data was analyzed using the ΔΔCt method with GAPDH as the housekeeping gene and D0 cells expression as a control. Box plots diagrams shows IQR (box) ± 1.5 outliers (whiskers) and mean (x). * p<0.05, ** p<0.01, **** p<0.0001, N.S. no statistical significance.

**Figure 6.**
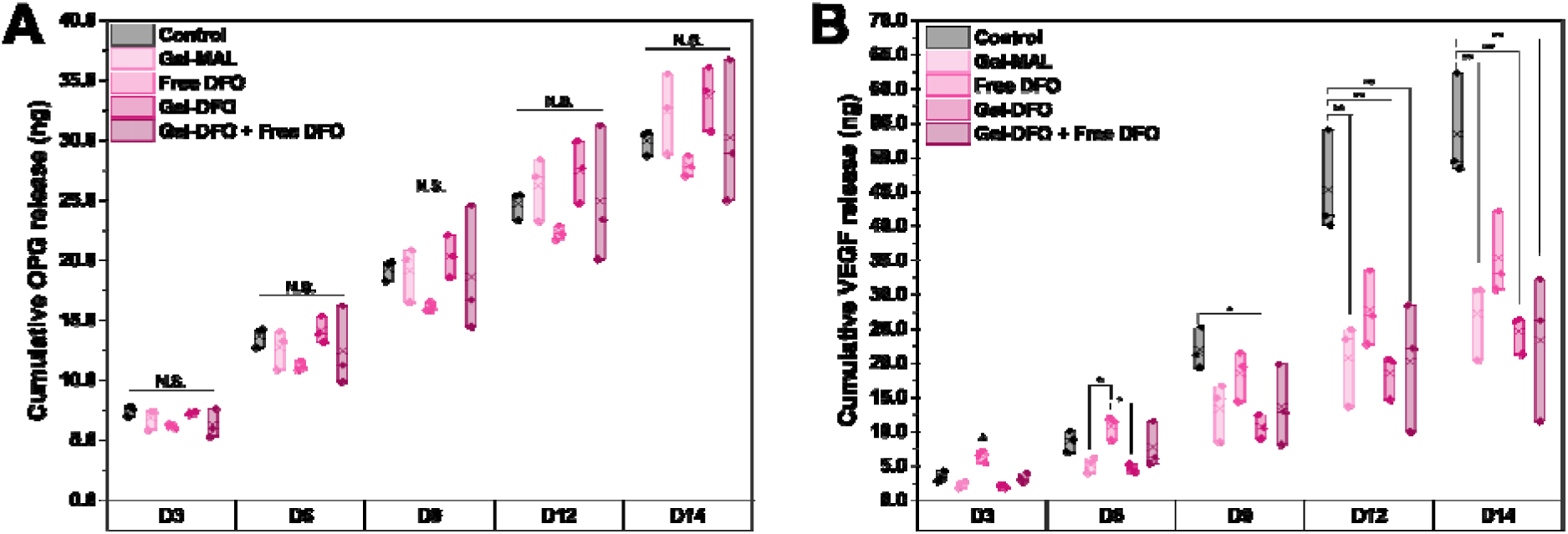
Secretome analysis. **A**. Cumulative OPG secretion. **B**. Cumulative VEGF secretion. Box plots diagrams show IQR (box) ± 1.5 outliers (whiskers) and mean (x). * p<0.05, ** p<0.01, N.S. means no statistical significance. Special character (^) represents that group has statistical significance (p<0.0001) compared to all groups on day 3.

### 3.6. Conditioned media from DFO-functionalized scaffolds induces endothelial cell network formation

In addition to considering a single angiogenic factor (VEGF), we also tested the overall angiogenic activity of conditioned media generated by hMSCs within mineralized collagen scaffolds with or without the addition of soluble DFO or DFO-functionalized GelMAL hydrogels. Conditioned media generated after 3 or 14 days in culture was collected and then applied to human umbilical vein endothelial cells (HUVECs) seeded on a Matrigel^TM^ assay plate. Resultant endothelial networks formed after 12 h of culture were analyzed via the Angiogenesis Analyzer Macros (ImageJ; Fig. 7A). Overall, the presence of soluble or hydrogel bound DFO in mineralized collagen scaffolds produced conditioned media after 3 days that significantly increased the complexity (number of junctions) of the resulting endothelial networks (Fig. 7B). We did not observe any effect of DFO condition on the length of individual endothelial network branches, with the average branch length formed measuring approximately 75 µm (Fig. 7C).

**Figure 7.**
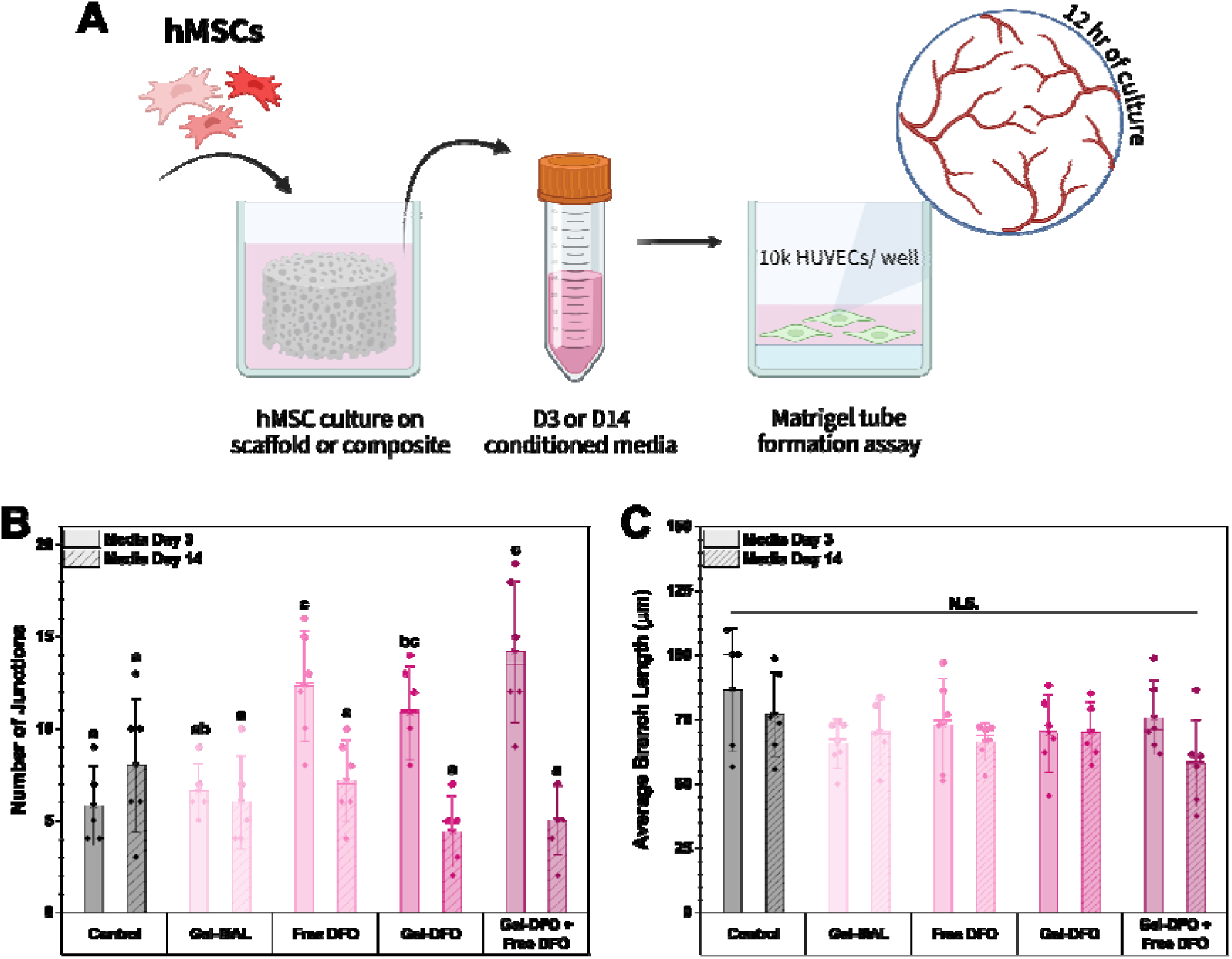
Tube formation assay. **A**. Experimental setup for Matrigel tube formation assay. Day 3 or 14 conditioned media from composite DFO-Collagen experiment was pooled and transferred into a Matrigel pre-coated plate with 10,000 HUVECs per well. Cells were cultured for 12 hours before brightfield images were taken on a Leica DMI8 confocal microscope. Images were analyzed with ImageJ Angiogenesis Analysis Macros. **B**. Number of junctions formed after 12 hrs. **C.** Average branch length after 12 hrs. Bar plots represent average value ± SD. Different letters denote statistical differences (p<0.05) and N.S. means no statistical difference.

## 4. Discussion

Angiogenesis is essential for wound healing and bone regeneration, as blood vessels transports nutrients and oxygen, as well as recruits cells into the injury area [70, 71]. However, the design requirements to promote osteogenesis vs. angiogenesis complicate material fabrication. Optimal angiogenesis has been shown in larger (~400 µm) pores, while osteogenesis in smaller pore structures (100 - 200 µm) [72–74]. Incorporation of activity-inducing biomolecules may help address this challenge by boosting the angiogenic potential of bone-specific biomaterials. Multiple avenues exist to incorporate angiogenic growth factors or drugs into biomaterials, but these often require supraphysiological doses, leading to hypertrophic tissue formation or rapid biomolecule clearance from the bloodstream or injury site [38, 39, 47]. Here, we explored the effects of incorporating an angiogenic supplement (DFO) tethered to a gelatin macromer into a class of mineralized collagen scaffold to form a multi-material (scaffold-hydrogel) composite to boost hMSC angiogenic and osteogenic activity.

We first identified DFO concentrations necessary to promote hMSC angiogenic activity without causing significant cytotoxicity. hMSCs were directly seeded on the mineralized collagen scaffold, and DFO was introduced into the cell media on experimental day 0. Increasing DFO concentration reduced hMSC proliferation but activated hypoxic pathways beneficial for wound healing and angiogenesis, findings consistent with prior study of DFO [44, 46]. Gene expression analyses confirmed DFO regulates angiogenic genes expression of HIF-1α and VEGF. Although changes were significant at early culture stages (p = 7.37e-7 and 1.48 e-4 for HIF-1a and VEGF at day 3, respectively), they were also transient due to the nature of soluble DFO exposure. Nonetheless, 20 µM soluble DFO was sufficient to maintain cell proliferation rates similar to the control, reduce hypoxic stress levels by day 14, and achieve VEGF expression levels similar to those at higher DFO concentrations on day 3. With prior work showing prolonged DFO exposure accelerated wound healing and bone formation [38], we hypothesized retaining DFO in the mineralized collagen scaffold via an implanted hydrogel carrier could maintain long-term DFO bioavailability.

Prior work showed DFO could be bound via its primary free amine to carboxylic acid groups within hyaluronic acid or to proteins via photo crosslinking [38, 75]. Although similar chemistry could have been applied to the collagen scaffold, we focused on approaches that would not affect the composition and crosslinking of the mineralized collagen scaffold already optimized for osteogenic activity. Rather, we used carbodiimide chemistry to attach DFO directly to a gelatin macromolecule functionalized with maleimides. While we anticipated a reaction connecting COOH groups on gelatin with amines on DFO, NMR and IR spectra suggest that DFO is also bound to GelMAL macromers via the maleimide group (31% decrease in maleimide degree of functionalization). Notably, we showed via a FeCl_3_ chelation assay that the vast majority of DFO was tethered to GelMAL hydrogels with no detectable DFO released after 24 h and the GelMAL bound DFO readily available for release (e.g., via hydrogel degradation) and formation of a chelation complex. Mixing DFO-functionalized and non-functionalized GelMAL macromers, we were able to create a variety of DFO dosed hydrogels to evaluate the effect of bound DFO on encapsulated hMSC activity. Similar to previous data with soluble DFO, hMSC proliferation was reduced initially in response culture within DFO-laden hydrogels; however, groups under 200 µM DFO showed no significant difference compared to conventional GelMAL hydrogels after day 3. VEGF secretion was increased with DFO over time, confirming that DFO-laden GelMAL hydrogels promote hMSC VEGF secretion in a dose-dependent manner up to an optimal concentration (200 µM). The 200 µM GelMAL-DFO concentration was identified for further composite models, as it released more VEGF compared to the 0 µM DFO control group. The results of the soluble and hydrogel DFO delivery assays were consistent with prior studies of DFO that have identified the need to balance the angiogenic vs cytotoxic effects of increasing concentration of DFO [38, 45, 76].

We then evaluated a multi-material composite approach that combined DFO-laden GelMAL hydrogels with the bolus addition of soluble DFO to hMSC seeded mineralized collagen scaffolds. Here, rapid exposure to soluble DFO could promote early angiogenic activity but would be likely limited with the DFO-laden hydrogels, motivating a model of bolus-release (soluble DFO) and sustained-availability (GelMAL-DFO hydrogel). As expected, DFO reduced cell proliferation at early culture time points, but cells recovered after the first week with a 4-fold increase for scaffolds containing both soluble and hydrogel-bound DFO (Gel-DFO + Free DFO). We evaluated genomic and functional consequences of DFO exposure via established markers of angiogenic (VEGF) and osteogenic (OPG) activity [22, 59, 77]. Osteoprotegerin (OPG) is a marker for osteoclast inhibition and has been shown in previous studies to regulate bone resorption and reduce osteoclast activity in our mineralized collagen scaffolds. We wanted to ensure this characteristic was preserved in our model, as osteoclastogenesis is essential for bone healing. OPG secretion profiles did not change with the inclusion of DFO or a hydrogel matrix, suggesting that the collagen scaffold still retains osteogenic potency. The observed reduction of VEGF secretion by hMSCs in response to DFO was not expected but suggests future opportunities for study. The reduced VEGF production at early culture times could have been due to reduced overall number of cells. At later stages when cell numbers were similar, reduced VEGF in the media could have been a result in their depletion in the media during the increased cell proliferation seen in DFO groups. Future studies measuring VEGF secretion from cell lysate rather than cell media would be needed to support this explanation.

Results from our composite approach also offer suggestions for future optimization of multi-material biomaterial to control both burst and sustained-release biomolecule availability. The observed upregulation of angiogenic genes in DFO-functionalized groups compared with control groups suggests opportunities to control the temporal availability of factors. Notably, while HIF-1α expression fluctuated for the control (scaffold alone, scaffold plus GelMAL) and soluble DFO groups, groups containing DFO-laden hydrogels showed steady HIF-1α upregulation (0.86 – 1.15 fold change). This plateau in the expression suggests sustained DFO induced HIF-1α stabilization, avoiding its degradation in the presence of oxygen. The Gel-DFO + Free DFO group shows a clear peak in VEGF expression on day 3. While longer sustained VEGF upregulation would be desirable, persistent VEGF upregulation may also produce detrimental effects similar to those observed with supraphysiological doses [78], potentially impairing healthy bone formation. Interestingly, while we observed HIF-1α stabilization for the Gel-DFO group, this did not translate into a VEGF peak like the Gel-DFO + Free DFO group. This suggests that early DFO exposure may be sufficient to boost VEGF upregulation, but extended DFO exposure may be necessary to maintain HIF-1α stabilization [79–81].

Lastly, knowing that single factors (e.g., VEGF) may not be sufficient to capture the complexity of the angiogenic activity of hMSC laden collagen biomaterials, we used conditioned media from the DFO-functionalization experiments to perform a Matrigel assay that measured the formation of HUVECs networks. Prior literature showed increase vascular formation with increasing DFO concentration [38]. Here, we explored whether the early (3 day) or late (14 day) secretome of hMSCs in response to mineralized collagen scaffolds and soluble and/or GelMAL hydrogel bound DFO would induce significant changes in HUVEC network formation. While no differences in branch length were observed, we observed an increased number of junctions in response to secretome from DFO-containing scaffolds at day 3, with no effect in late-stage media (day 14). This suggests that DFO drove early changes in hMSC secretome that promoted new vessel sprouting. While the Matrigel assay is a good first generation model to assess the effect of shifting hMSC secretome on angiogenesis, future work could like use more complex in vitro angiogenic assays that examine tube formation as well as in vivo implantation or chick embryo chorioallantoic membrane (CAM) assays to more deeply examine the role of DFO on scaffold angiogenesis. Regardless, biomaterials that promote both osteogenic and angiogenic processes are urgently needed to aid the regeneration of functional bone tissue. This work established methods to incorporate a pro-angiogenic small molecule into porous materials using a tunable secondary hydrogel phase that could assist angiogenic potency without disrupting the primary matrix osteogenic properties.

## 5. Conclusions

Vascular integration is necessary for a successful biomaterial bone graft. While we have previously described scaffolds that promote osteogenesis, cues for simultaneous angiogenic processes remain limited. This work proposes development of multi-material approaches to act as both osteogenic and angiogenic promoters by combining porous collagen scaffolds with DFO-laden hydrogel to balance DFO release and resultant activity of human mesenchymal stem cells. This work identified a soluble DFO concentration window that supports hMSC viability and upregulation of angiogenic activity within mineralized collagen scaffolds, both in soluble and hydrogel bound forms. We also report a strategy to chemically tether DFO to a gelatin macromer with 79.6% retention over 7 days while maintaining its iron chelation activity. Lastly, we describe a multi-material composite design featuring porous mineralized collagen scaffolds containing DFO-laden gelatin hydrogels. This novel construct shows that DFO-functionalized scaffolds promote stabilized HIF-1α expression while the addition of soluble DFO upregulated early-stage VEGF expression and promoted endothelial network branching. Together, this work expands our ability to temporally retain deferoxamine at physiological levels, and with a practical fabrication strategy tailored to osteogenic and angiogenic coupling mechanisms.

## Acknowledgements

The authors would like to acknowledge undergraduate students Zoyia Khan^1^, Tara Mukherjee^2^ and research scientist Alison C. Nunes^2^ (^1^Department of Bioengineering and ^2^Chemical and Biomolecular Engineering; University of Illinois Urbana-Champaign) for their assistance with data collection. The authors would also like to acknowledge the following institutes: the Carl R. Woese Institute for Genomic Biology, the Beckman Institute for Advanced Science and Technology, School of Chemical Sciences Mass Spectrometry Laboratory, and the Roy J Carver Biotechnology Center, all at the University of Illinois Urbana-Champaign. Gene expression analyses were performed at the Roy J. Carver Biotechnology Center Shared Resource at the Cancer Center at Illinois (CCIL), which is partially supported by the NCI Cancer Center Support Grant (CCSG) award number P30CA275774. Research reported in this publication was supported by the National Institute of Arthritis and Musculoskeletal and Skin Diseases of the National Institutes of Health under Award Numbers R01 AR077858 (BACH) and R01 AR078892 as well as the National Institute of Dental and Craniofacial Research of the National Institutes of Health under Award Number R01 DE030491 (BACH). We are grateful for the funding for this study provided by the Grainger College of Engineering SURGE Fellowship (GRA) and Sloan UCEM Scholarship (GRA). The content is solely the responsibility of the authors and does not necessarily represent the official views of the National Institutes of Health. This manuscript is the result of funding in whole or in part by the National Institutes of Health (NIH). It is subject to the NIH Public Access Policy. Through acceptance of this federal funding, NIH has been given a right to make this manuscript publicly available in PubMed Central upon the Official Date of Publication, as defined by NIH.

## Contributions (CRediT: Contributor Roles Taxonomy [82, 83])

**G. Ríos Adorno:** Conceptualization, Data curation, Formal Analysis, Validation, Visualization, Investigation, Methodology, Writing – original draft, Writing – review & editing. **A. Bale:** Data curation, Investigation, Validation, Writing – review & editing. **C. Subramanian:** Conceptualization, Methodology, Writing – review & editing. **R. Bhargava**: Resources, Writing – review & editing. **M.S. Cohen:** Conceptualization, Methodology, Writing – review & editing. **B.A.C. Harley:** Conceptualization, Resources, Project administration, Funding acquisition, Supervision, Writing – review & editing.

