## Supplemental Information for "Multi-material delivery strategies to create deferoxamine-functionalized collagen scaffolds for craniofacial bone repair"

^a^ Dept. Chemical and Biomolecular Engineering

^b^ Carle Illinois College of Medicine

^c^ Carl R. Woese Institute for Genomic Biology

^d^ Cancer Center at Illinois

^e^ Dept. of Bioengineering

^f^ Beckman Institute for Advanced Science and Technology

University of Illinois Urbana-Champaign

Urbana, IL 61801, USA

**Corresponding Author:**

B.A.C. Harley

Dept. of Chemical and Biomolecular Engineering

Cancer Center at Illinois

Carl R. Woese Institute for Genomic Biology

University of Illinois at Urbana-Champaign

110 Roger Adams Laboratory

600 S. Mathews Ave.

Urbana, IL 61801

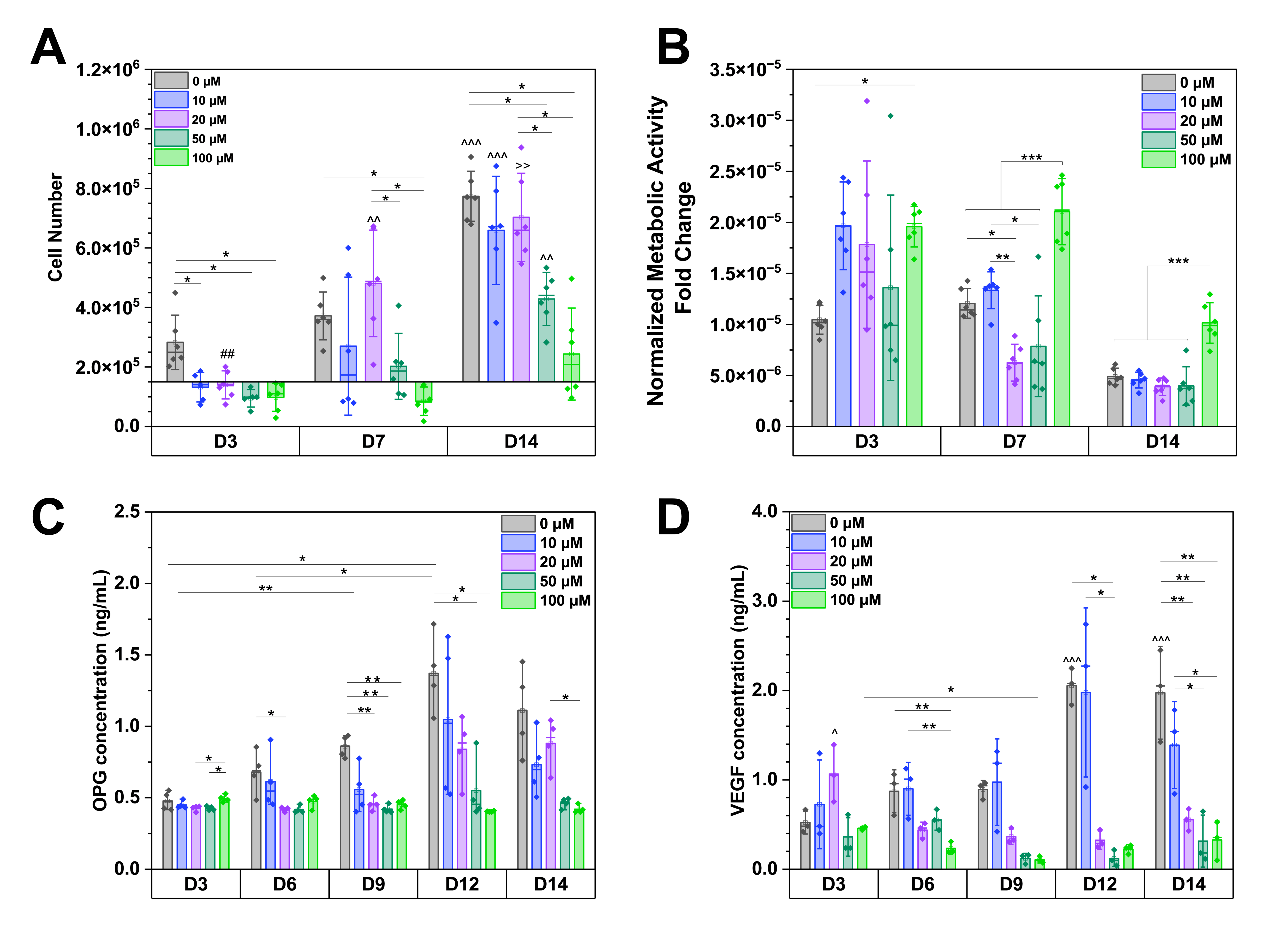

**Supplementary Figure S1. DFO supplemented media effects on hMSCs**. **A.** Cell number (n=6) across time points. **B.** Metabolic activity (n=6) normalized to group cell number at each time point. **C**. OPG (ng/mL, n=4) concentration across days of culture. **D**. VEGF (ng/mL, n=3) concentration across days of culture. Bar plot diagrams show the mean ± SD. * p<0.05, ** p<0.01 and *** p <0.001. Special characters represent statistical significance in the same concentration at different time points.

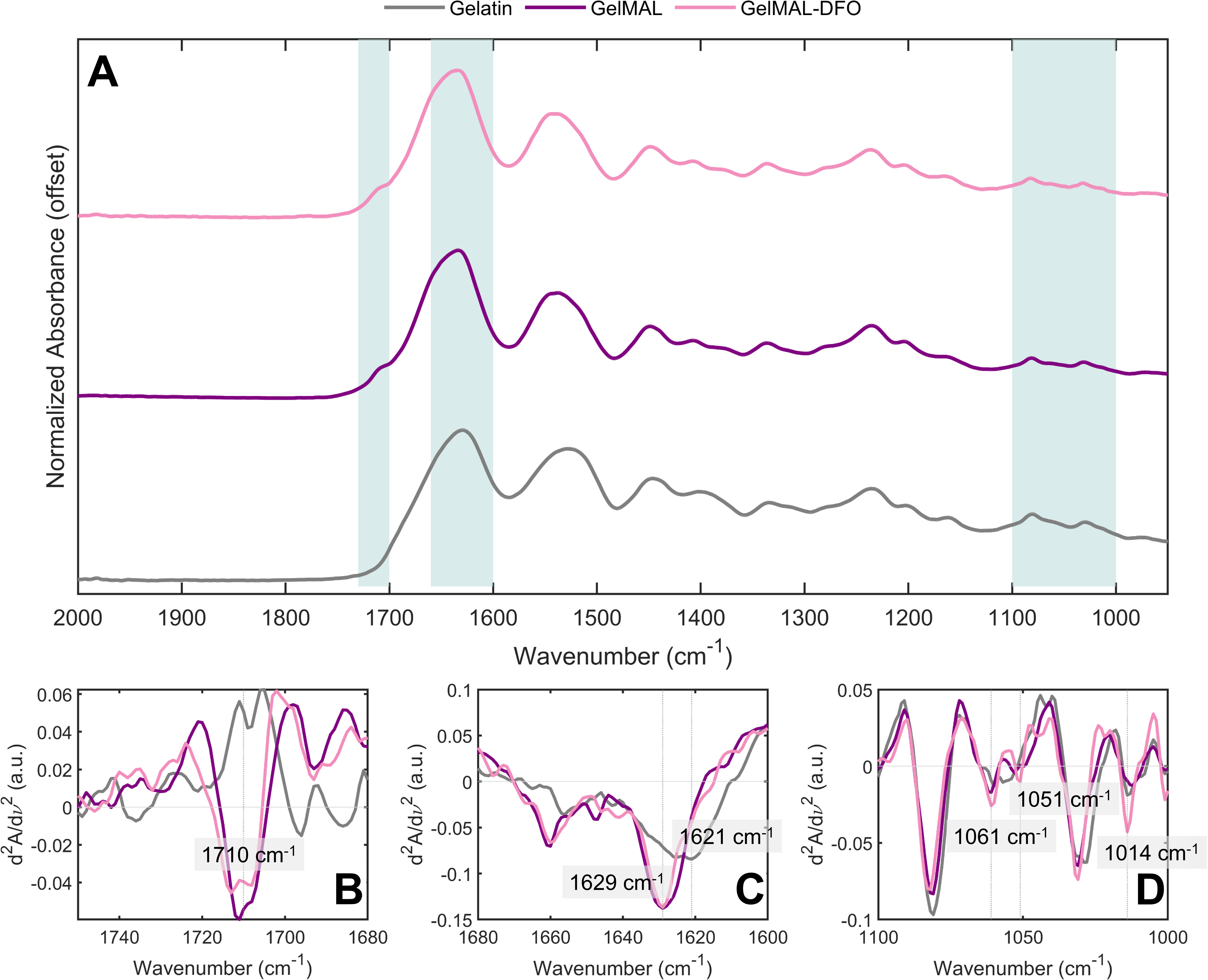

**Supplementary Figure S2. IR spectra of gelatin materials acquired in attenuated total reflectance using an FT-IR spectrometer.** **A.** Spectra 2000-950 cm^-1^. **B.** Carbonyl stretching vibrations of the maleimide group (1700 - 1750 cm^-1^). **C.** Amide region I, carbonyl stretching vibration of the peptide backbone (1621 cm^-1^ – gelatin, 1629 cm^-1^ – GelMAL and GelMAL-DFO). **D.** Hydroxamate N-O stretching of DFO (1000-1100 cm^-1^).
